# *Lactiplantibacillus plantarum* Membrane Vesicles (MVs) Exhibit Immunomodulatory and Bactericidal Effects Against *Escherichia coli* and *Salmonella* Typhimurium

**DOI:** 10.1101/2025.08.26.672515

**Authors:** Cristal Dafne Lonngi Sosa, Francisco Rodolfo González Díaz, Hugo Ramírez Álvarez, Alejandro Vargas Ruíz, José Luis Muciño Hernández, Rosa Isabel Higuera Piedrahita, Héctor Alejandro de la Cruz Cruz, Marisela Leal Hernández, Gerardo Ramírez-Rico, Jorge Alfredo Cuéllar Ordaz, Cynthia González Ruíz

## Abstract

The indiscriminate use of antibiotics has led to an increase in multidrug-resistant bacteria, necessitating the search for effective alternative therapies to reduce antimicrobial use. Lactic Acid Bacteria (LAB) have been explored as an alternative to antibiotics due to their multiple beneficial properties. These bacteria secrete membrane vesicles (MVs), key acellular components in combating pathogens. This study aimed to evaluate the effects of *Lactiplantibacillus plantarum* MVs co-cultured with *Escherichia coli* (MVsplE) or *Salmonella* Typhimurium (MVsplS) through inhibition assays using disk diffusion on agar plates, as well as their activation and cytokine expression in the RAW 264.7 macrophage cell line. The results showed that MVsplE and MVsplS were produced in greater quantity and size than non-co-cultured *L. plantarum* MVs (MVspl). Additionally, MVsplE and MVsplS exhibited a dose-dependent inhibitory effect on the growth of enteropathogenic bacteria. Furthermore, RAW 264.7 cells stimulated with these MVs demonstrated that the expression of IL-1β, TNF-α, and IL-10 depended on the enteropathogenic strain with which *L. plantarum* was previously co-cultured. Following a challenge with enteropathogenic bacteria, the MVs induced an immunomodulatory response. These findings demonstrate that *L. plantarum* MVs exert bactericidal and immunomodulatory effects against enteropathogenic bacteria, suggesting their potential use as an alternative treatment to antimicrobials.

**Author summary:** Lactic acid bacteria have been widely used as probiotics in therapies aimed at controlling or treating infectious diseases. However, it has been reported that these bacteria can replicate in immunocompromised individuals, as well as in clinically healthy individuals, potentially disrupting the intestinal microbiota and contributing to disease progression. In recent years, bacterial membrane vesicles (MVs) have emerged as promising alternatives, demonstrating superior effects compared to whole bacteria. These MVs are nanostructures derived from the cytoplasmic membrane and are released at various stages of bacterial growth. They contain multiple biomolecules and structural membrane proteins that enable them to exert biological effects on the host. Notably, MVs do not replicate, and therefore do not disturb the intestinal microbiota, leading to their classification as acellular probiotics. Research on these biological agents is crucial to minimizing risks to host health and developing alternative treatments for gastrointestinal diseases. In this study for first time, we demonstrated the bactericidal and immunomodulatory effects of *Lactiplantibacillus plantarum* MVs against *Escherichia coli* and *Salmonella* Typhimurium, supporting their potential as an effective alternative in the treatment of gastrointestinal infections.

## Introduction

Gastrointestinal tract diseases are often caused by imbalances in the microbiota of animals, which can lead to health issues by weakening the immune system and acting as a risk factor for disease development **[1, 2]**. The pathogens involved in these processes pose a significant public health concern, as they are transmitted through contaminated food, water, or direct contact with infected animals, resulting in gastroenteric diseases in humans **[3]**. Among these pathogens, *Escherichia coli* is one of the most critical zoonotic agents; in 2024 was responsible for approximately 265,000 infections, 3,600 hospitalizations, and 30 deaths annually in the United States. Similarly, *Salmonella* species reported in 2003 approximately 93.8 million cases of gastroenteritis and 155,000 deaths annually. To address this issue, new preventive strategies have been implemented, primarily aimed at reducing transmission through contaminated animal-derived products, thereby decreasing the number of infected animals and, consequently, the number of human cases **[3, 4]**. The use of probiotics in animal diets has been explored due to their demonstrated antimicrobial efficacy, their positive impact on animal production, and their safety in terms of public and environmental health **[5]**. The most used genera include *Lactococcus* spp.*, Streptococcus* spp*., Leuconostoc* spp.*, Pediococcus* spp.*, Lactobacillus* spp.*, Bifidobacterium* spp.*, Weissella* spp., and *Lactiplantibacillus plantarum* (formerly *Lactobacillus plantarum*), which belong to the Lactic Acid Bacteria (LAB) group LAB are Gram-positive bacteria naturally present in the gastrointestinal microbiota and possess immunomodulatory, antimicrobial, and pro-inflammatory properties, as well as the ability to enhance mucus production at the intestinal level **[6–8]**.

The beneficial properties of LAB are primarily attributed to their cellular structure and the release of soluble antigens into the environment. Recent studies have highlighted the crucial role of MVs, which are secreted at any stage of bacterial growth **[9]**. MVs are spherical structures ranging from 40 nm to 400 nm in diameter **[10]**. In Gram-positive bacteria, MVs originate from the cytoplasmic membrane and carry colonization factors, bactericidal components, DNA, RNA, and possible remnants of the bacterial cell wall **[11–15]**. In addition to these components, MVs possess a selective charge determined by bacterial growth conditions and derived from their formation. These structures allow bacteria to create favorable niches for replication and colonization **[16–18]**. The large number of antigens contained in MVs can interact with and trigger responses in innate immune system cells through the expression of different interleukins **[19]**. Studies have reported that *L. plantarum* MVs can induce the expression of both pro-inflammatory and anti-inflammatory cytokines in cell cultures, including IL-6, TNF-α, IL-1β, IL-10, IFN-γ, and IL-12 **[12, 20, 21]**. However, these studies have exclusively employed ATCC strains of *L. plantarum*. Additionally, proteomic analyses have demonstrated the antimicrobial and probiotic effects of *L. acidophilus* MVs, which contain ABC transporters and bacteriocins, potential key components of these effects **[22]**.

The objective of this study was to evaluate the effect of *L. plantarum* MVs co-cultured with *Escherichia coli* or *Salmonella* Typhimurium, as determined through bacterial inhibition assays, and their impact on a murine macrophage cell line (RAW 264.7). Our findings revealed that *L. plantarum* MVs co-cultured with *Escherichia coli* (MVsplE) or *Salmonella* Typhimurium (MVsplS) exhibited a greater concentration and size, as well as a dose-dependent bactericidal effect against the evaluated enterobacteria. Moreover, MVsplE and MVsplS promoted RAW 264.7 activation and primarily stimulated IL-10 expression, exerting an immunomodulatory effect upon challenge with *Escherichia coli* or *Salmonella* Typhimurium.

## Materials and methods

### Bacterial Strains

For this study, a strain of *Lactiplantibacillus plantarum* was isolated and purified from the colon of clinically healthy rats captured in urban settlements of Mexico City. These rats underwent a quarantine period, followed by euthanasia and necropsy following NOM-062-ZOO-1999 and the protocol approved by the by the Institutional Subcommittee for the Care and Use of Experimental Animals (SICUAE) of the Facultad de Estudios Superiores Cuautitlán, UNAM, under approval number MC-2022/1-1, dated March 27, 2022. The strain was molecularly characterized using endpoint PCR and biochemically identified with the Apiweb 50CHL system (BioMériux, Lyon, France). *Lactobacillus acidophilus* ATCC 314 (Thermo Scientific, Massachusetts, USA) strain was used as a positive control in the PCR assays. Field strain of *Escherichia coli* and a strain of *Salmonella enterica* serovar Typhimurium ATCC 154 were donated by Instituto Nacional de Investigaciones Forestales, Agrícolas y Pecuarias (CENID-INIFAP, Mexico City, Mexico) for co-culture assays with *L. plantarum*.

### Molecular Characterization

For molecular characterization, DNA was extracted using cetyltrimethylammonium bromide (CTAB) (Sigma-Aldrich, Massachusetts, USA) and the phenol-chloroform method **[23]** from different cultures isolated from the gastrointestinal tract (GIT) of free-living rats. To identify lactic acid bacteria (LAB) belonging to the genera *Lactobacillus* spp., *Leuconostoc* spp., *Pediococcus* spp., and *Weissella* spp., the following primers were used: Forward: 5’AGCAGTAGGGAATCTTCCA 3’, Reverse: 5’ATTYCACCGCTACACATG 3’. To specifically identify *Lactobacillus* spp., the following primers were used: Forward: 5’TGGATGCCTTGGCACTAGGA 3’, Reverse: 5’AAATCTCCGGATCAAAGCTTACTTAT 3’. A *Lactobacillus acidophilus* ATCC 314 strain was used as a positive control, and molecular grade water was used as a negative control. The PCR amplification conditions were as follows: Initial denaturation: 5 min at 95°C, 35 cycles of: Denaturation: 30 s at 95°C, o Annealing: 30 s at 56°C (for LAB) and 62°C (for *Lactobacillus* spp.), extension: 30 s at 72°C, final extension: 10 min at 72°C. Amplification products were visualized on 2% agarose gels stained with Midori Green (Nippon Genetics, Tokyo, Japan), using the Orange G ruler 50 bp (Thermo Scientific, Massachusetts, USA) molecular weight marker as a reference.

### Co-culture and Challenge Experiments

For co-culture and challenge experiments, a field strain of *Escherichia coli* and a strain of *Salmonella enterica* serovar Typhimurium ATCC 154 were used. To evaluate the antimicrobial effect, *L. plantarum* was co-cultured with enteropathogenic bacteria following the methodology of Savino et al., 2011, with modifications. Tryptic Soy Broth (TSB) (Dibico, State of Mexico, Mexico) and Lactobacilli MRS (BD, New Jersey, USA) media were used. Briefly, a 35 µL aliquot of *L. plantarum* (1×10^6^, 0.05 OD) grown in Lactobacilli MRS medium was incubated for 24 h at 37°C under 5% CO₂ in static conditions. After that, it was cultured in 35 mL of TSB for five h (3×10^6^, 0.15 OD) at 37°C under 5% CO₂ in static conditions, aliquots of 100 µL were taken periodically to ensure the pH remained above 5, thereby confirming that bactericidal activity was associated with bacterial metabolic products. Subsequently, *E. coli* (1×10⁸ CFU) or *S*. Typhimurium (1×10^8^ CFU) was added, and co-cultures were incubated for 24 h at 37°C **[24]**.

A negative control was included, in which enteropathogenic bacteria were grown in MRS and TSB media under the same conditions as *L. plantarum* to verify that the inhibitory effect was due to the presence of LAB and not by the medium change. After incubation, Gram staining was performed, and light microscopy confirmed bacterial inhibition. The absence of Gram-negative bacteria was considered a positive result for *L. plantarum* mediated inhibition. Additionally, to confirm the inhibition, co-cultures with *E. coli* or *S*. Typhimurium were cultivated on MacConkey agar (BD, New Jersey, USA). The absence of growth in these media confirmed the inhibitory effect of *L. plantarum.* Experiments were conducted in triplicate with independent samples.

Complete cells (C.C) obtained from *L. plantarum* co-cultured with enteropathogenic bacteria were designated as “C.CplE” when co-cultured with *E. coli* and “C.CplS” when co-cultured with *S*. Typhimurium. These samples were later used to obtain membrane vesicles (MVs) for subsequent assays.

### Isolation and Quantification of *L. plantarum* MVs

*L. plantarum* strains previously co-cultured with *E. coli* or *S*. Typhimurium were centrifuged at 1,400 × g for 3 min. The supernatant was discarded, and the pellet was resuspended in 500 mL of Lactobacilli MRS medium and incubated aerobically for 24 h at 37°C to induce stress and enhance MVs production. Cultures were then centrifuged at 9,000 × g for 15 min, and the supernatant was sequentially filtered through nitrocellulose membranes with 0.45 µm and 0.22 µm pore sizes before ultracentrifugation at 150,000 × g for three h at 4°C. The resulting pellet, corresponding to the MVs fraction, was resuspended in 500 µL of sterile PBS and stored at −80°C until use **[25]**.

Protein quantification was performed using the Bradford method with linear regression and a bovine serum albumin (BSA) standard curve. Experiments were conducted in triplicate with independent samples **[26]**.

### Transmission Electron Microscopy of MVs

*L. plantarum* samples and MVs were placed on 200-mesh copper grids coated with formvar (Electron Microscopy Sciences, Pensilvania, USA) and shadowed with carbon to confirm MV formation and assess purification. A 10 µL sample was applied to the grid and stained with 1% phosphotungstic acid (pH 6.0) (Sigma-Aldrich, Massachusetts, USA) for 1 min. Samples were visualized using a JEM 1400 (JEOL, Peabody, Massachusetts, USA) transmission electron microscope at the Research and Advanced Studies Center of the National Polytechnic Institute (CINVESTAV), Zacatenco Unit **[27]**.

### Characterization of MVs by NanoSight NS300

To evaluate MVs size and concentration, samples were ultrafiltered through 0.22 µm membranes. MVs number and distribution were analyzed by NanoSight NS300 (Malvern Panalytical, Malvern, UK) [28], at the National School of Biological Sciences, Santo Tomás Unit, National Polytechnic Institute (IPN).

### Antimicrobial Inhibition Assays of MVsplE or MVsplS from *Lactiplantibacillus plantarum* on Enteropathogenic Bacterial Cultures

After confirming the formation and purification of MVsplE or MVsplS, antimicrobial inhibition assays were performed. The methodology of Vanegas et al. (2017) was followed with some modifications. Petri dishes were prepared with a base layer of Mueller-Hinton agar (Dibico, State of Mexico, Mexico) and allowed to solidify at room temperature. Subsequently, a layer of previously sterilized and tempered Sulfide, Indole, Motility (SIM) semi-solid medium (BD, New Jersey, USA) was added, containing *Salmonella* Typhimurium (1×10^8^) or *Escherichia coli* (1×10⁸). The plates were then refrigerated at 4°C for 2 hours [29].

Once solidified, sensi-disks were placed with the following treatments: 75 or 100 µg of MVspl, MVsplE, or MVsplS, respectively. Experimental controls included C.Cpl, C.CplS, C.CplE, or sterile PBS. After applying the treatments at different concentrations, the culture plates were incubated for 18–24 hours at 37°C. These assays were performed in triplicate with independent samples.

### Stimulation of RAW 264.7 Cells with *L. plantarum* MVs and Challenge with Enteropathogenic Bacteria

To evaluate the stimulation and possible activation of macrophages with *L. plantarum* MVs, the methodology of Gutiérrez et al. (2023) was followed with some modifications. The RAW 264.7 murine macrophage cell line (ATCC, Virginia, USA) was cultured in 24-well plates for 8 hours at a concentration of 1×10⁵ cells per well in high-glucose DMEM medium (4.5 g/L) (Biowest, Nuaillé, France) supplemented with 10% fetal bovine serum (HyClone, Utah, USA). The cells were incubated for 24 hours at 37°C with 5% CO₂ to ensure adherence to the culture plates. RAW 264.7 cells were stimulated by adding 10 µg of MVs or C.C from *L. plantarum*, either co-cultured with enteropathogenic bacteria or non-co-cultured. Additionally, 2 µg of lipopolysaccharide (LPS) (*Escherichia coli* O111:B4, Sigma-Aldrich, Massachusetts, USA) or 1× PBS were used as experimental controls [25].

Throughout the 8-hour experiment, cells were stimulated at 0, 2, and 4 hours. At 5 hours, they were challenged with 10 µL of a dilution containing *E. coli* (1×10⁸) and *S.* Typhimurium (1×10^8^), respectively. The cultures were then incubated for an additional 3 hours to reach a total of 8 hours. Samples were collected at 0, 1, 3, 5, and 8 hours for qPCR analysis, performing three independent experimental replicates.

### qPCR of IL-1β, TNFα, and IL-10 in RAW 264.7 Cells Stimulated with MVs and Challenged with Enteropathogenic Bacteria

To determine interleukin expression, RNA was extracted from cultured cells at 0, 1, 3, 5, and 8 hours using TRIzol Reagent (Thermo Scientific, Massachusetts, USA). RNA was purified using the chloroform-isopropanol-ethanol method [25]. The resulting precipitate was resuspended in 50 µL of RNase-free water. RNA concentration was measured by spectrophotometry using a NanoDrop (NanoDrop Lite, Thermo Scientific, Massachusetts, USA). Subsequently, cDNA was synthesized using the FastGene Scriptase Basic cDNA Synthesis Kit (Nippon Genetics, Tokyo, Japan). The concentration of the obtained cDNA was determined by spectrophotometry. Primers for IL-1β, TNFα, and IL-10 were synthesized by T4Oligo (Irapuato, Guanajuato, Mexico) based on published sequences from GenBank (IL-1β #NM_008361.4, TNFα #NM_001278601.1, and IL-10 #NM_010548.2). Primers were designed using Primer3 (v.0.4.0) and aligned using BioEdit software (BioEdit v7.2.5, Ibis Bioscience, California, USA). The sequences are listed in Table 1. qPCR was performed in triplicate with independent samples for each interleukin, using 10 ng of cDNA and a Master Mix (RealQ Plus Master Mix Green Without ROX, AMPLIQON, Odense, Denmark). Amplification was carried out using an Agilent Technologies Mx3005P system (Stratagene Mx3000P, Thermo Scientific, Massachusetts, USA).

**Table 1.**
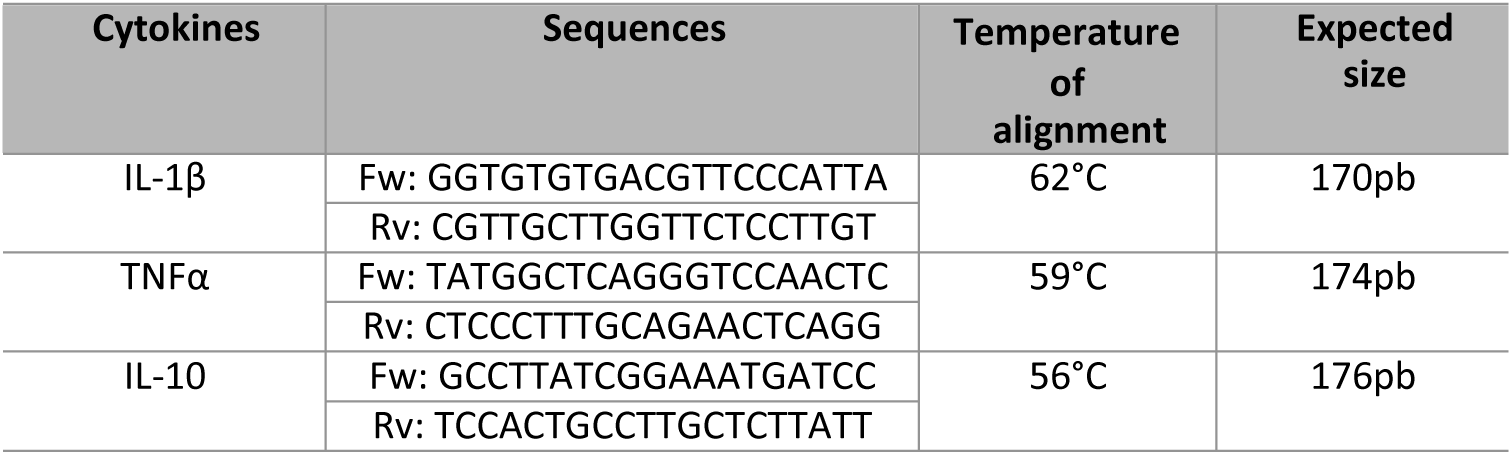
Primer Sequences and Annealing Parameters for IL-1β, TNFα, and IL-10.

The housekeeping gene hypoxanthine-guanine phosphoribosyltransferase (HPRT) was used as an internal control. The amplification protocol consisted of the following steps: Enzyme activation: 1 cycle at 95°C for 15 min, Denaturation: 95°C for 30 s, Annealing: 50°C for 30 s, Elongation: 72°C for 30 s (40 cycles). The amplification conditions for cytokines were identical, except for the annealing temperature, which is detailed in **Table 1**. Amplification and dissociation curves were generated to verify the specific expression of the gene of interest (S1 Fig).

Relative expression quantification was performed using the ΔΔCt method, applying the following equations:

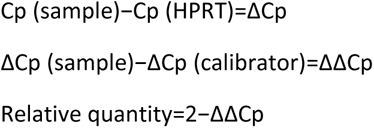

Subsequently, the logarithmic transformation of the relative values obtained was performed using the base 10 logarithm ([2–ΔΔCp (Log 10)]) **[30, 31]**.

### Statistical Analysis

Data were analyzed using a one-tailed Student’s t-test and analysis of variance (ANOVA) followed by Tukey’s test. Statistical analyses were conducted using GraphPad Prism 8.0.2 (GraphPad, California, USA). Differences were considered significant when *p* < 0.1 (*), *p* < 0.05 (**), and *p* < 0.001 (***).

## Results

### Identification and Molecular Characterization of *Lactiplantibacillus plantarum*

Following the isolation of bacterial cultures from free-living rats, molecular characterization was performed using PCR to identify the LAB genus in which the evaluated strain showed a single band of 350 bp, coinciding with the expected product size (Fig 1A). Once confirmed, a second PCR was conducted to determine the *Lactobacillus* genus; the strain evaluated showed a single band of 100 bp, consistent with the expected product size (Fig 1B). Finally, biochemical identification was carried out using the Apiweb 50CHL system, where the strain exhibited 99.4% identity with *Lactobacillus plantarum*, which has recently been reclassified as *Lactiplantibacillus plantarum* **[7]**.

**Fig 1.**
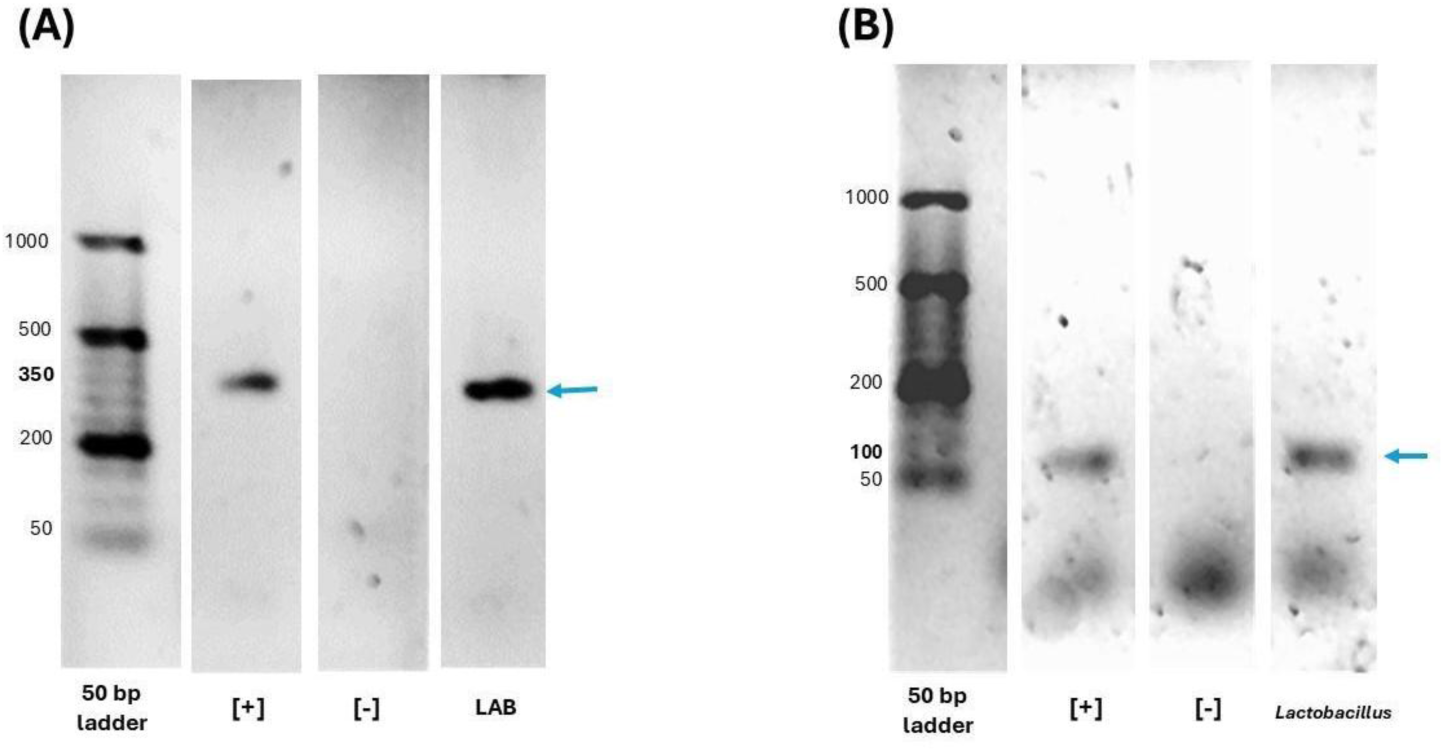
Molecular and biochemical identification of *Lactiplantibacillus plantarum.* Molecular identification was performed by PCR on 2% agarose gels stained with Midori Green, using the Orange G ruler 50 bp molecular weight marker. *L. acidophilus* ATCC 314 strain was used as a positive control (+), and molecular-grade water served as the negative control (−). (A) Electrophoresis Gel corresponding to the LAB genus, showing a single 350 bp band. (B) Electrophoresis Gel for the *Lactobacillus* genus (previously *Lactobacillus plantarum*), showing a single 100 bp band. All samples were from a single gel.

### Bactericidal Effect of *Lactiplantibacillus plantarum* Against *E. coli* and *S*. Typhimurium

Subsequently, the strain of *L. plantarum* was identified, and its effect on *E. coli* and *S.* Typhimurium was evaluated through co-culture experiments. Our results demonstrated that *L. plantarum* inhibited the growth of enteropathogenic bacteria in TSB and MRS media. Gram-staining and cultures on agar plates, where only enteropathogenic bacteria grew, were performed to verify this inhibition. Microscopy analysis revealed the absence of Gram-negative bacteria (S2 Fig); similarly, agar plates did not show growth of *E. coli* or *S.* Typhimurium (Fig 2A and 2B) when co-cultures of *L. plantarum* with *S.* Typhimurium or *E. coli* were performed, respectively. In contrast, in the negative control, where only enteropathogenic bacteria were grown, normal growth was observed in all media tested, confirming that the inhibition was due to *L. plantarum* and not to changes in the culture medium (Fig 2C and 2D).

**Fig 2.**
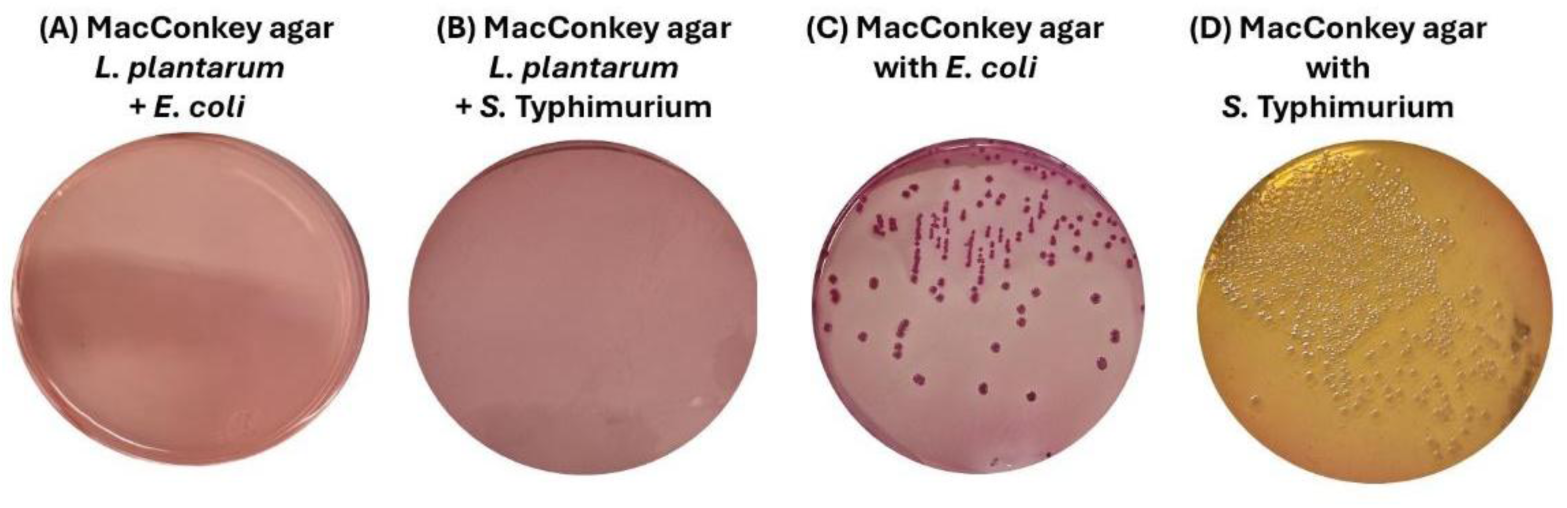
*L. plantarum* co-cultures viability on agar plates. (A) *E. coli* 1×10⁸ strain after being co-cultured with *L. plantarum* in MacConkey agar. (B) *S.* Typhimurium 1×10⁸ strain after being co-cultured with *L. plantarum* in MacConkey agar. (C) *E. coli* strain after being cultured in TSB and MRS in MacConkey agar. (D) *S*. Typhimurium strain after being cultured in TSB and MRS in MacConkey agar. The absence of growth of *E. coli* or *S*. Typhimurium in their specific media after being co-cultured with *L. plantarum*, demonstrated the bactericidal effect of *L. plantarum*.

### Co-Culture of *L. plantarum* with Enteropathogenic Bacteria Stimulates MVs Secretion

After confirming the antimicrobial effect of *L. plantarum* in co-cultures with enteropathogenic bacteria, we isolated MVs to assess their secretion and characterization using TEM. Negative staining TEM analysis of *L. plantarum* MVs co-cultured with *Salmonella* Typhimurium (MVsplS) or *Escherichia coli* (MVsplE) revealed the formation of multiple MVs originating from the cytoplasmic membrane (Fig 3A and 3B), along with their aggregation on the peptidoglycan layer (Fig 3C). TEM also confirmed the purification of MVs, which displayed a spherical shape, a diameter ranging from 100 to 200 nm, and the ability to coalesce (Fig 3D and 3E). Compared to non-co-cultured *L. plantarum* MVs (MVspl), fewer vesicles were observed (Fig 3F).

**Fig 3.**
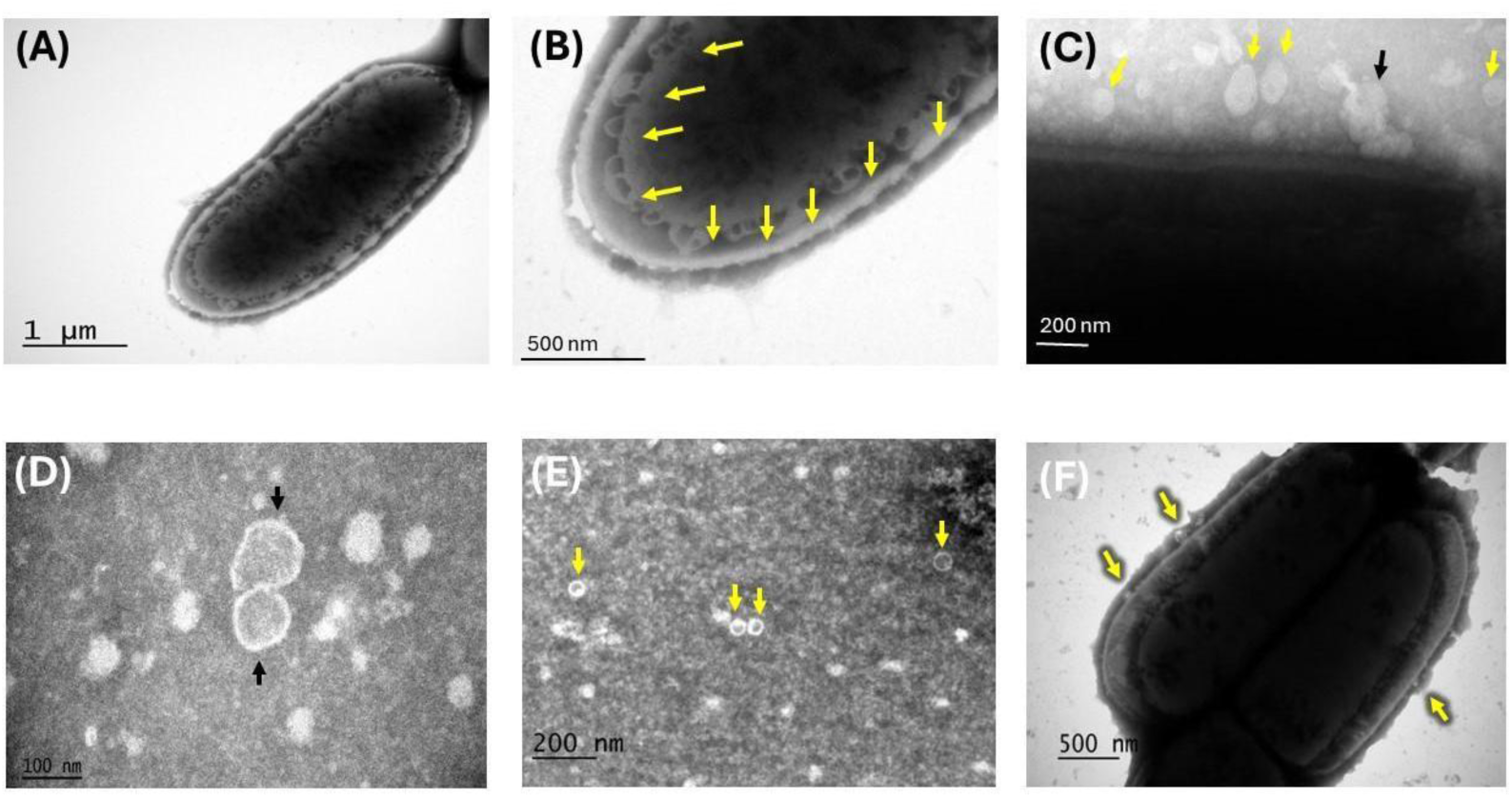
Negative-stained transmission electron microscopy of *L. plantarum* MVs. (A) The formation of multiple MVsplS can be seen along the cytoplasmic membrane of the bacterial cell. (B) Close-up of the cytoplasmic membrane where the yellow arrows show the formation of multiple MVs. (C) MVsplS (yellow arrows), the black arrow indicates their possible fusion. (D) MVsplE with a spherical shape, an approximate diameter of 100 nm, and their coalescence capacity. (E) Multiple MVsplE are shown in this panel (yellow arrows). (F) Formation of MVspl (yellow arrows) was found in smaller quantities compared to those co-cultured with enteropathogenic bacteria.

### Co-Culture of *L. plantarum* with *E. coli* and *S*. Typhimurium Increases MV Production and Size

Given that TEM revealed a higher number of MVsplE and MVsplS compared to MVspl, we further evaluated their concentration and size using the NanoSight NS300 system. The results showed that MVsplE and MVsplS exhibited increased size and concentration compared to MVspl **(Table 2)**. These findings were corroborated by video recordings from the NanoSight NS300 (S1, S2 and S3 video).

**Table 2.**
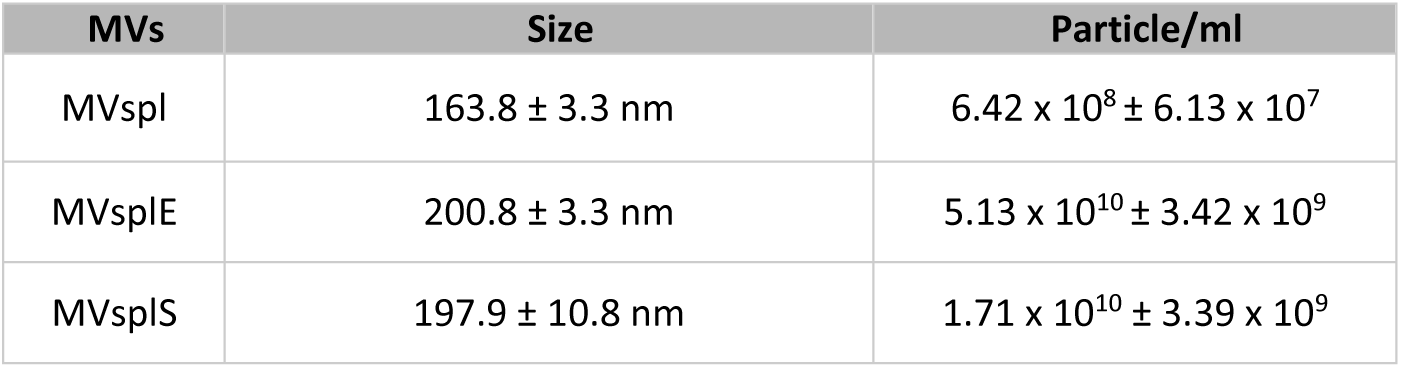
Differences in size and concentration between MVspl, MVsplE, and MVsplS.

| MVs | Size | Particle/ml |
| --- | --- | --- |
| MVspl | 163.8 ± 3.3 nm | 6.42 x 10<sup>8</sup> ± 6.13 x 10<sup>7</sup> |
| MVsplE | 200.8 ± 3.3 nm | 5.13 x 10<sup>10</sup> ± 3.42 x 10<sup>9</sup> |
| MVsplS | 197.9 ± 10.8 nm | 1.71 x 10<sup>10</sup> ± 3.39 x 10<sup>9</sup> |

### Antimicrobial Effect of MVsplE and MVsplS Against Enteropathogenic Bacteria

Once MVsplE and MVsplS were isolated, antimicrobial inhibition assays were performed using disk diffusion tests. The size of the inhibition zones was dependent on the concentration of MVs and complete cells obtained from *L. plantarum* (C.Cpl), complete cells of *L. plantarum* co-cultured with *E. coli* (C.CplE) or complete cells of *L. plantarum* co-cultured with *S.* Typhimurium (C.CplS), inoculated on the disks, with a more substantial effect observed at 100 µg of MVs compared to their complete cells, respectively. In that sense, 100 µg of MVsplE showed a greater size inhibition compared to the negative control (*p* = 0.06) (Fig 4A), as did MVsplS (*p* = 0.03) (Fig 4B). Notably, 100 µg of MVsplS demonstrated the highest inhibitory effect against *S.* Typhimurium, forming a 2.5 cm inhibition zone (Fig 5). The inhibition zones of MVsplE and MVsplS exhibited irregular edges, possibly due to their smaller size and higher diffusion capacity in the medium **[32]**.

**Fig 4.**
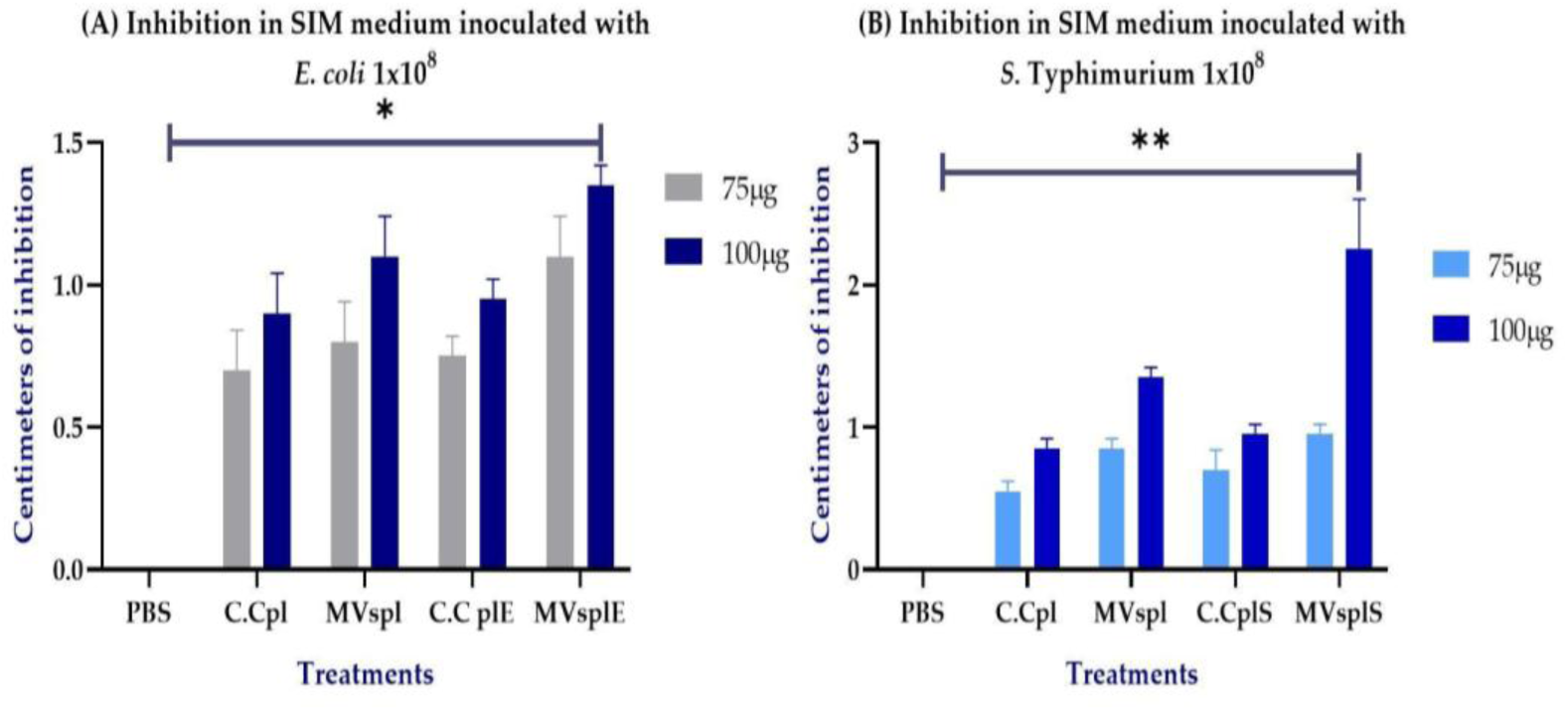
Inhibition of *E. coli* or *S*. Typhimurium bacteria by *L. plantarum* MVs in an agar disc diffusion test. (A) Inhibition in SIM medium inoculated with *E. coli* at 1×10^8^ with 75 and 100 µg of MVspl or MVsplE. (B) Inhibition in SIM medium inoculated with *S*. Typhimurium 1×10^8^ with 75 and 100 µg of MVspl or MVsplS. Analysis performed by one-way ANOVA, results represent \**p* < 0.1 and \*\**p*< 0.05. As positive control, C.Cpl, C.CplE, or C.CplS was used, while PBS served as the negative control. Analysis was performed using one-way ANOVA; results are represented as \**p* < 0.1 and \*\**p* < 0.05.

**Fig 5.**
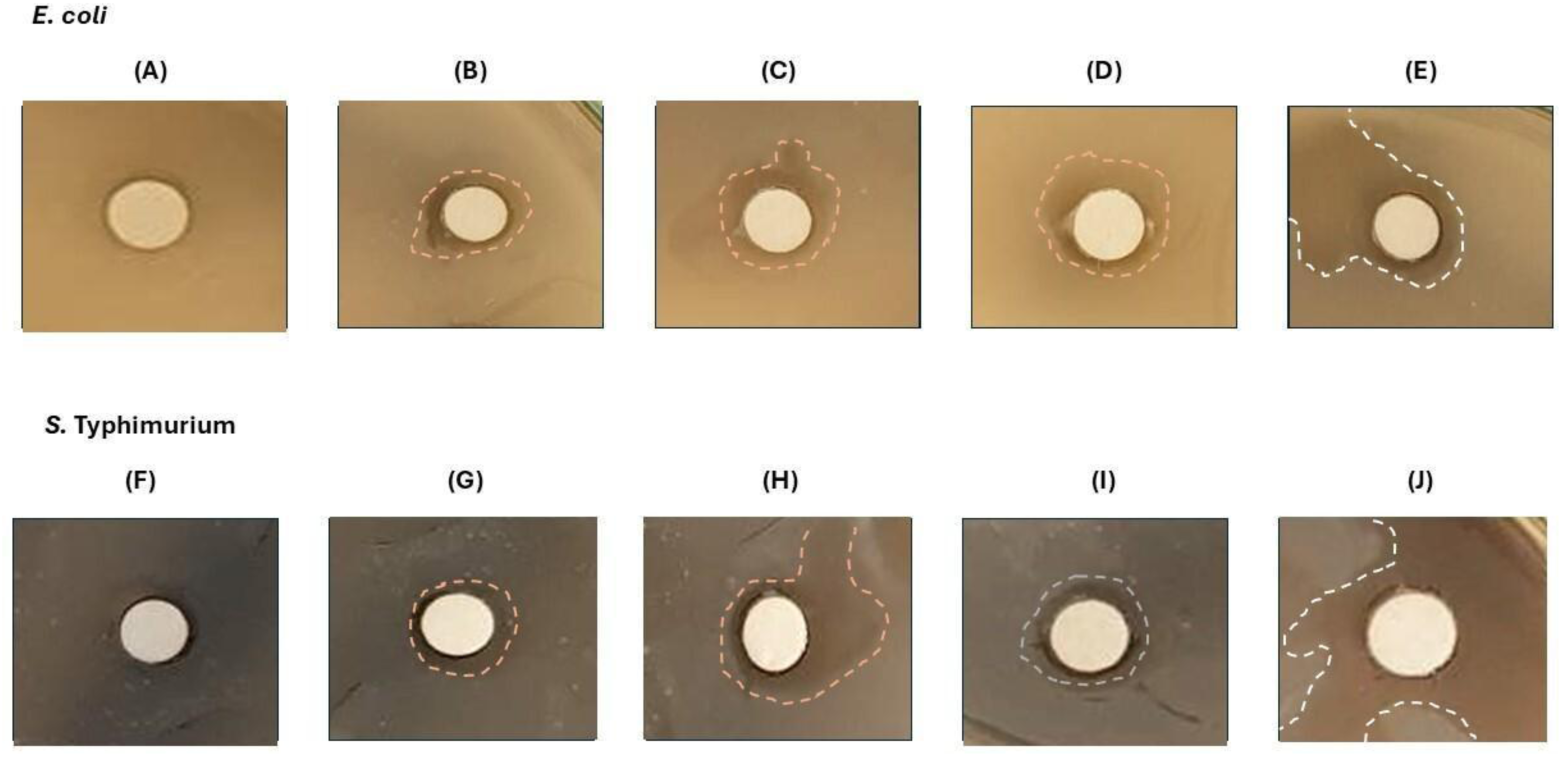
Inhibition zone in SIM medium inoculated with enteropathogenic bacteria, with the addition of 100 µg of the different treatments. SIM medium inoculated with *E. coli* 1×10^8^. (A) PBS, (B) C.Cpl, (C) MVspl, (D) C.CplE, and (E) MVsplE. In addition, the SIM medium was also inoculated with *S.* Typhimurium 1×10^8^ CFU/mL, (F) PBS, (G) C.C60pl, (H) MVspl, (I) C.CplS, and (J) MVsplS.

### Administration of MVsplE or MVsplS Triggers Activation of RAW264.7 Cells

Macrophages are professional antigen-presenting cells present in the host’s mucus (including the digestive system). They are also one of the primary cells involved in inflammatory responses through the secretion of cytokines. Following antigenic processing, these cells actively participate in the inflammatory process, either promoting or inhibiting the response, and polarizing toward the M1 or M2 profile, respectively **[33, 34]**. To evaluate the stimulation and potential activation of Raw 264.7 cells with MVs, the macrophages were treated with various conditions (Fig 6). Before the experiments, the RAW 264.7 macrophage cell line was cultured in 24-well plates for 24 hours to ensure adherence. Previously, the cells exhibited spherical and regular morphology under stimulation (Fig 6A). However, after treatment, they adopted a spindle-shaped morphology with pseudopodia (Fig 6B-H). Although no significant morphological differences were observed between the different treatments, we proceeded to assess the expression of pro-and anti-inflammatory cytokines.

**Fig 6.**
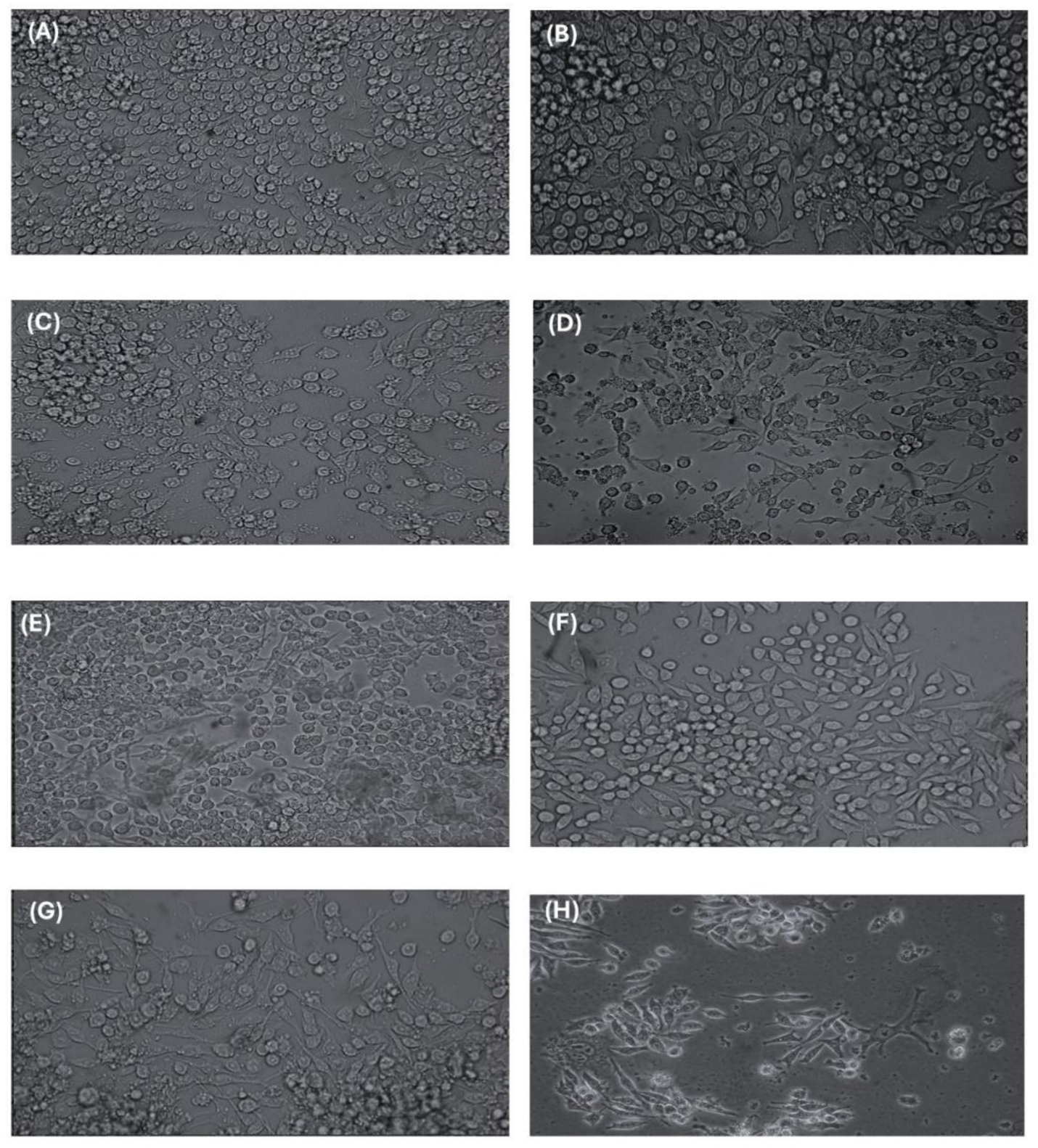
Morphological changes observed in RAW 264.7 at hour 5 after being stimulated at hour 0, 2 and 4. (A) Cells with PBS, (B) Cells incubated with two µg of LPS, (C) Cells stimulated with C.Cpl, (D) Cells stimulated with MVspl, (E) Cells incubated with C.CplE, (F) Cells incubated with MVsplE, (G) Cells with C.CplS, (H) Cells with MVsplS. All groups treated with C.C. or MVs were stimulated with 10µg, respectively.

### Cytokine Expression in RAW264.7 Cells Is Modulated by MVsplE and MVsplS

Following the observed morphological changes in RAW264.7 cells after MVs administration, the expression of cytokines IL-1β, TNF-α, and IL-10 was evaluated using quantitative PCR (qPCR). Macrophages stimulated with MVsplE or MVsplS displayed a pro-inflammatory profile characterized by increased IL-1β and TNF-α expression from the beginning to hour 3 of the experiment. However, macrophages stimulated with MVsplE from hour 5 until the end of the challenge (hour 8) exhibited higher IL-10 expression than IL-1β (*p* = 0.01) and TNF-α (*p* = 0.09) (Fig 7A).

**Fig 7.**
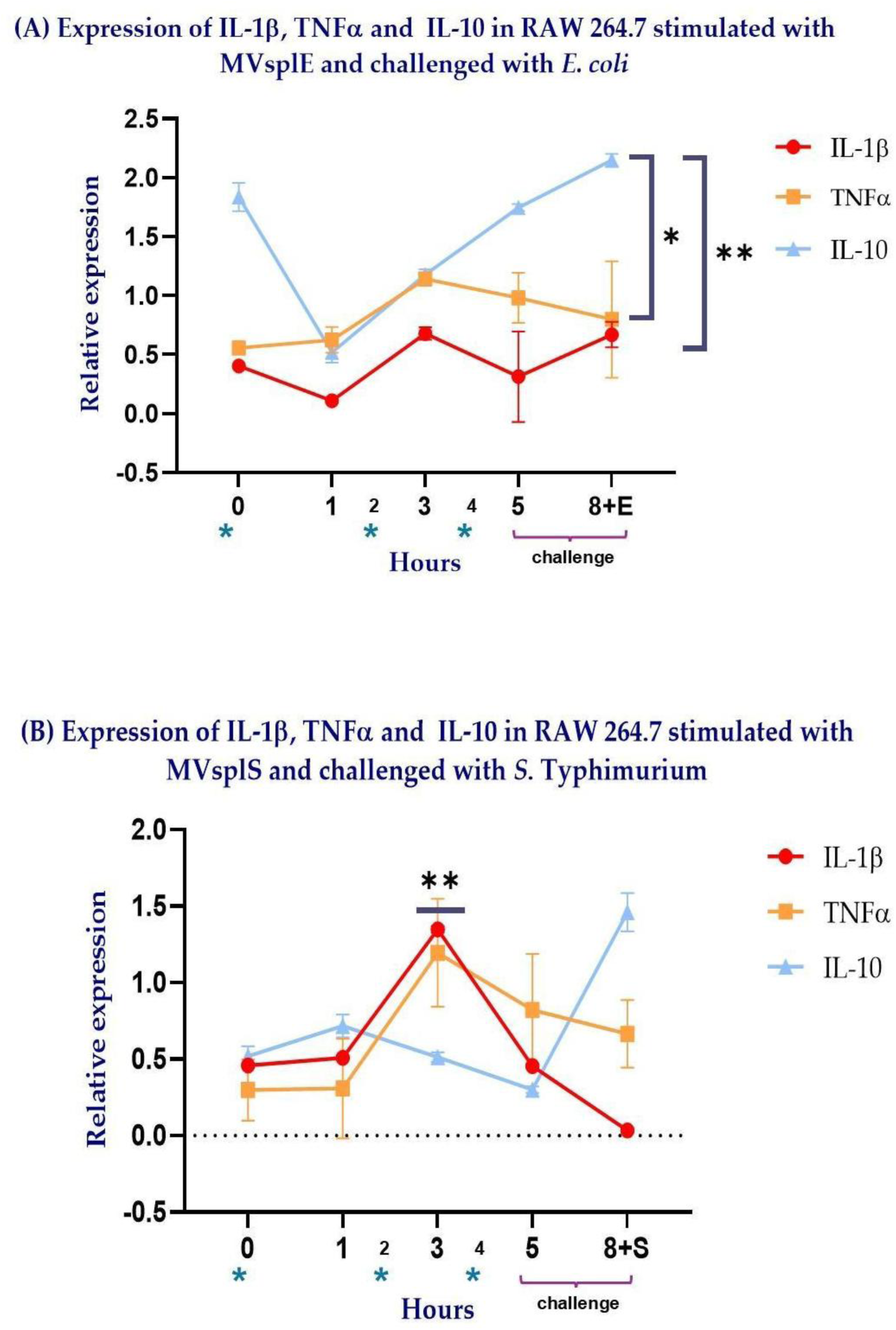
Kinetics of expression of IL-1β, TNF-α, and IL-10 in RAW 264.7 cells, which were stimulated at hours 0, 2, and 4, respectively (*), with 10µg of the different MVs. Subsequently, RAW 264.7 cells were challenged with enteropathogenic bacteria at hour 5 and finished at hour 8. (A) Expression of cytokines in RAW 264.7 stimulated with MVsplE and challenged with *E. coli*. (B) Expression of cytokines in RAW 264.7 stimulated with MVsplS and challenged with *S*. Typhi-murium. Analysis performed using Student’s t-test; results are represented as \**p* < 0.1 and \*\**p* < 0.05.

In contrast, macrophages stimulated with MVsplS showed greater expression of IL-1β at hour 3 compared to IL-10 (*p = 0.005).* After this period, at hour 5, the expression of IL-1β and IL-10 exhibited a sharp decrease. Subsequently, at hour 8, following the S. Typhimurium challenge, TNF-α and IL-10 were expressed, with the latter exhibiting its highest expression level (Fig 7B).

Based on these results, the next step was to compare cytokine expression in macrophages between the different groups. To do this, the individual expression of the different cytokines was analyzed.

### RAW264.7 Stimulation with MVsplE or MVsplS Induces Higher Cytokine Expression Compared to LPS, C.C., and MVspl

Following kinetic analysis of cytokine expression in RAW264.7 cells stimulated with MVsplE or MVsplS, the expression of each cytokine was compared between the different groups. Regarding IL-1β expression, macrophages stimulated with MVs reached their peak expression at hour 3. Macrophages stimulated with MVsplE exhibited increased IL-1β expression compared to the negative control (*p* = 0.0028) (Fig 8A). Similarly, macrophages stimulated with MVsplS showed significantly higher IL-1β expression than the negative control (*p = 0.03*). Furthermore, at hour 3, it was observed that significantly higher IL-1β expression compared to the negative control (*p* = 0.0053) and the LPS control (*p* = 0.006) (Fig 8B). Although there was no statistically significant difference concerning MVs and their C.C regarding IL-1β expression, we observed that the expression in macrophages stimulated with MVsplS or MVsplE promoted a rapid and sustained IL-1β response, peaking at hour 3 and persisting until hour 5. In contrast, C.C-treated cells exhibited lower cytokine expression (Fig 8A and 8B).

**Fig 8.**
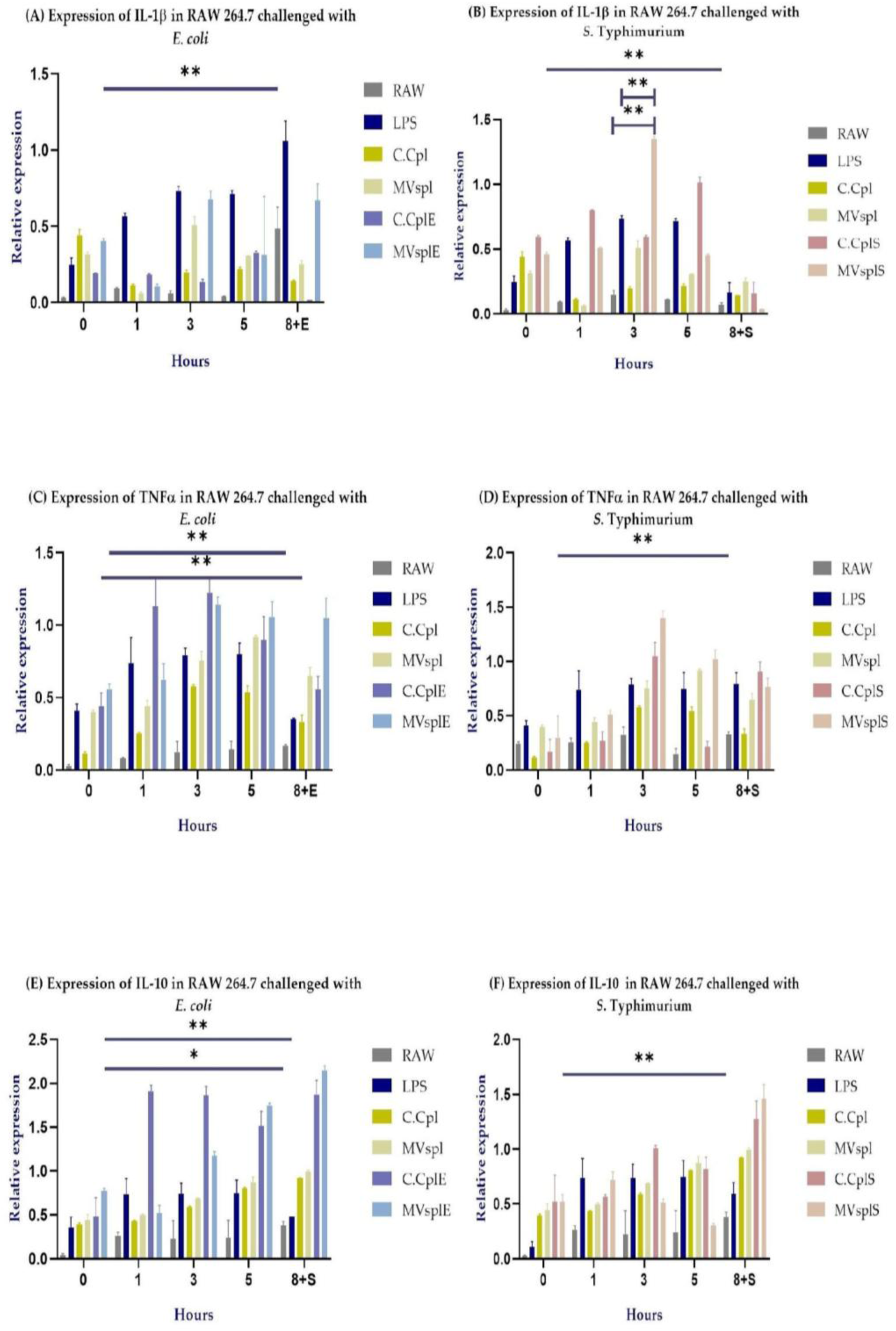
Cytokine expression in RAW 264.7 cells was stimulated at hours 0, 2, and 4 with 10 µg of MVs or C.C., or two µg of LPS, followed by a challenge at hour 5 with enteropathogenic bacteria, and concluded at hour 8. (A) Expression of IL-1β, (C) Expression of TNF-α, and (E) Expression of IL-10 in RAW 264.7 cells stimulated with MVsplE and challenged with *E. coli*. (B) Expression of IL-1β, (D) Expression of TNFα, (F) Expression of IL-10 in RAW 264.7 cells stimulated with MVsplS and challenged with *S.* Typhimurium. Analysis performed by two-way ANOVA; results represent \**p* < 0.1 and \*\**p* < 0.05.

Similarly, TNF-α expression reached its peak at hour 3 in macrophages stimulated with MVs. Macrophages stimulated with MVsplE or MVsplS exhibited higher expression compared to their respective negative controls (*p* = 0.0016 and 0.047, respectively) (Fig 8C and 8D). Likewise, the C.CplE showed a higher expression of this cytokine compared to the C.Cpl (*p* = 0.0157) (Fig 8C). While macrophages stimulated with MVsplS exhibited a similar response to IL-1β, they did not show a statistically significant difference compared to C.CplS. However, despite this, macrophages stimulated with MVsplS displayed a more rapid and sustained cytokine response compared to those treated with C.CplS (Fig 8D).

In RAW 264.7 cells stimulated with MVsplE, a higher expression of IL-10 was observed compared to the negative control (*p = 0.08)* and to macrophages stimulated with LPS *(p = 0.02*) (Fig 8E). Likewise, cells stimulated with MVsplS showed a higher expression of this cytokine than the negative control (*p* = 0.0490) (Fig 8F). In addition, we observed an interesting behavior: macrophages stimulated with MVsplE or MVsplS reached peak IL-10 expression at the end of the challenge with enteropathogenic bacteria (Fig 8E and 8F). This led us to question whether this effect was due solely to the action of MVs or if the presence of enteropathogenic bacteria was the trigger. For this reason, the next step was to evaluate cytokine expression in both challenged and unchallenged macrophages exposed to enteropathogenic bacteria.

### MVsplE and MVsplS Exert an Immunomodulatory Effect in RAW264.7 Cells Challenged with *E. coli* and *S*. Typhimurium via IL-10 Expression

Once we determined that the MVs on RAW264.7 cells showed a greater expression of IL-10 at the end of the challenge with enteropathogenic bacteria, we evaluated the secretion of this cytokine between different treatments. Macrophages stimulated with MVsplE plus *E. coli* showed a higher expression of IL-10 compared to cells stimulated with *E. coli* alone (*p* = 0.007) and even those stimulated with LPS plus *E. coli* (*p* = 0.04) (Fig 9A). On the other hand, macrophages stimulated with MVsplS plus *S*. Typhimurium promote a higher expression of this cytokine compared to cells with *Salmonella* (*p* = 0.02), as well as with those that were stimulated with LPS plus *Salmonella* (*p* = 0.006) and even with those that were stimulated with MVsplS (*p* = 0.009) (Fig 9B), confirming its immunomodulatory effect against *E. coli* and *S*. Typhimurium.

**Fig 9.**
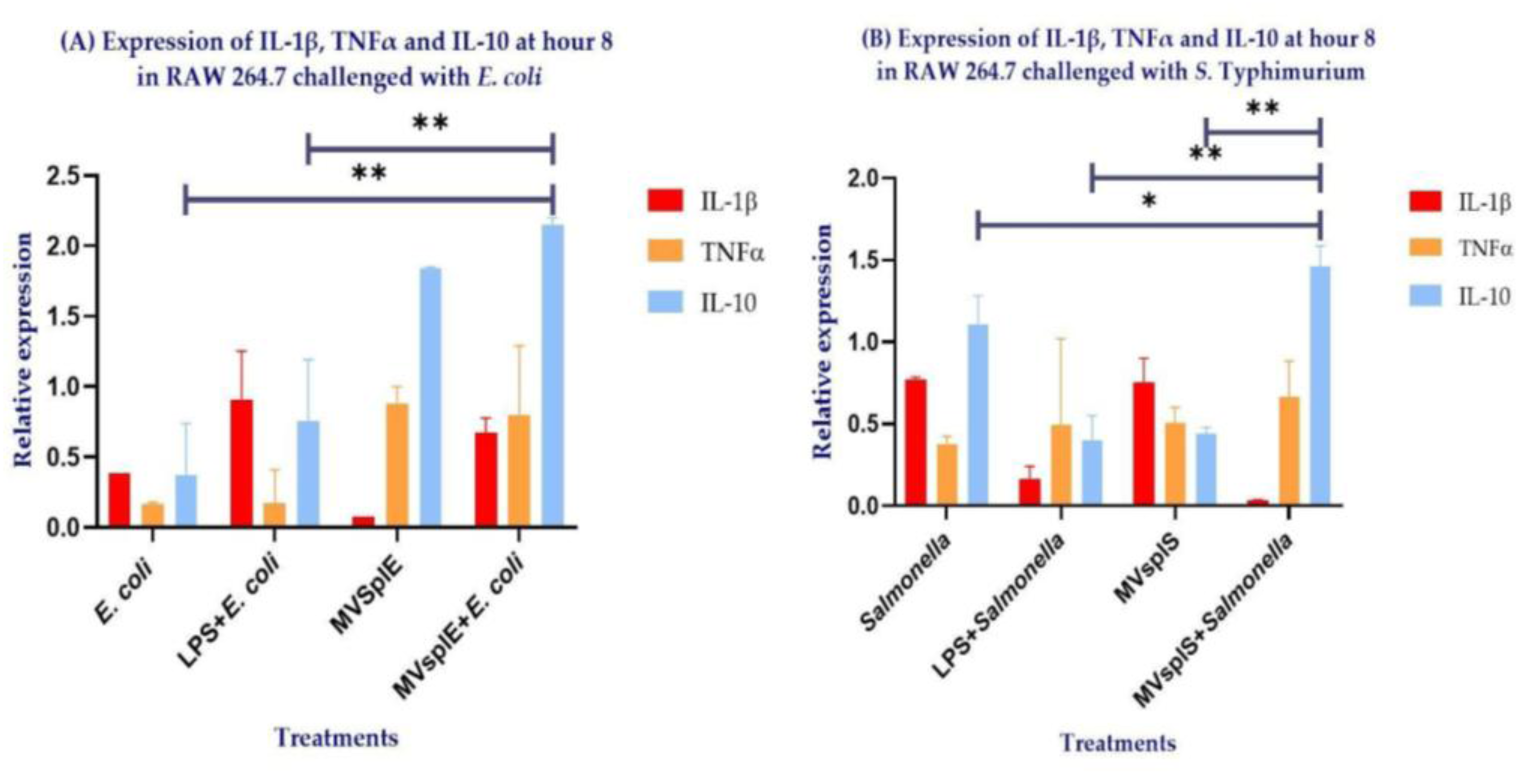
Cytokine expression at hour 8 in RAW 264.7 stimulated with (A) MVsplE and challenged with *E. coli* or (B) MVsplS and challenged with *S*. Typhimurium. All groups were stimulated at hours 0, 2, and 4 with their respective treatments, except for the RAW + *E. coli* and RAW + *Salmonella* groups, in which macrophages did not receive any stimulus. Likewise, all groups were challenged at hour 5 with the corresponding enteropathogenic bacteria, except for the MVsplE and MVsplS groups. Analysis performed by one-way ANOVA; results represent \**p* < 0.1 and \*\**p* < 0.05.

## Discussion

Antimicrobial resistance has increased recently, leading to a shortage of practical tools for preventing and treating drug-resistant infections. This has led to an increase in ineffective treatments [35]. Research in recent years has focused on finding alternatives that reduce the use of antibiotics. The properties of probiotics, such as immune system regulation and bactericidal capacity, play a crucial role in host health, even under certain pathological conditions. Therefore, the use of MVs from LAB has been proposed as a safe alternative to their original C.C. These structures lack replication capacity and significant transport of biologically active antigens as well as more efficient dispersion due to their size and structure, can quickly disperse through the thick mucus layer of the intestinal epithelium, allowing direct migration to other tissues and/or interaction with different immune cells in the gastrointestinal tract (GIT) compared to their C.C. counterparts [36].

In this study, we focused on MVs from LAB isolated from the GIT of wild rats residing in urban settlements, due to their ability to survive, proliferate, and reproduce in an environment with a high load of pathogenic microorganisms that are detrimental to other species, including humans. In 2015, Himosworth et al. demonstrated the prevalence of *E. coli* and *S.* Typhimurium in the GIT of *Rattus norvegicus* and *Rattus rattus*. Since the latter showed no signs or lesions of disease, this suggests that the LAB present in its GIT microbiota secretes potent antimicrobial agents and immunogenic components **[37].** In this study, we found that *L. plantarum* isolated from the colons of wild rats could inhibit the growth of *E. coli* and *S*. Typhimurium. These results align with findings by Shah et al. (2016), who demonstrated the antimicrobial effect of an ATCC strain of *L. plantarum* when co-cultured with *E. coli* for 24 hours. The authors attributed this effect to factors such as competition between *L. plantarum* and *E. coli* for nutrients in the culture medium, as well as the production of organic acids, hydrogen peroxide, fatty acids, and bacteriocins **[38]**. However, it is known that LAB secrete MVs into the medium at any growth phase, and these MVs carry bacteriocins and enzymes capable of exerting antimicrobial effects, as well as necessary adhesion, transport, and colonization antigens **[13]**. The characterization of MVsplE and MVsplS using TEM revealed their spherical shape and cytoplasmic membrane structure, along with the formation of multiple MVs at the cytoplasmic level, consistent with findings from various authors regarding Gram-positive bacteria and the *L. plantarum* genus **[18, 39]**. Additionally, Nanosight NS300 characterization of *L. plantarum* MVs, both co-cultured and non-co-cultured, showed that MVsplE and MVsplS had a larger size and higher concentration. This observation aligns with Orench-Rivera et al. (2016), who reported that the microenvironmental conditions to which bacteria are exposed strongly influence the secretion of MVs and the components carried within them. Co-cultured LAB with enteropathogenic bacteria exhibited increased MVs production, possibly carrying a more significant number of antigens *[16]*.

Furthermore, we confirmed the antimicrobial effect of LAB MVs on SIM medium inoculated with *S*. Typhimurium and *E. coli*, demonstrating that the antimicrobial effect is concentration-dependent. Larger inhibition halos (2.5 cm in diameter) were observed with 100 μg of protein compared to those obtained with 75 μg of protein (0.7 cm in diameter). These findings align with those of Lee (2021), who demonstrated the bactericidal effect of *L. plantarum* BCRC10357 MVs in agar diffusion assays against *Shewanella putrefaciens*, showing a bactericidal effect dependent on MV concentration **[28]**. However, this is the first study to demonstrate the antimicrobial impact of MVs from *L. plantarum* isolated from the colons of wild rats against *S*. Typhimurium and *E. coli*.

After confirming the antimicrobial effect of *L. plantarum* MVs, the murine macrophage cell line RAW 264.7 was stimulated with 10 μg of MVs, revealing morphological changes following different treatments with C.C and MVs from *L. plantarum* strains, both co-cultured and non-co-cultured. The macrophages transitioned from a spherical, regular shape to a fusiform shape with pseudopodia, possibly indicative of cellular activation. Xiaoyan (2022) observed similar morphological changes when stimulating RAW 264.7 cells with 10 ng/mL of commercial LPS, suggesting macrophage activation, which is consistent with our results **[40]**. Notably, microscopic analysis showed no morphological differences between the treatments, leading us to assess cytokine expression to identify potential polarization.

Macrophage activation is essential for responding appropriately to various stimuli. By altering their function in response to different microenvironmental signals, this process is known as polarization. Macrophages can polarize into two distinct profiles: M1 (classically activated) and M2 (alternatively activated). The M2 profile comprises subtypes, including M2a, M2b, M2c, and M2d, which are differentiated by their expression of specific markers, cytokines, and chemokines, thereby influencing their functions **[41]**. M1 macrophages are highly effective against intracellular pathogens and promote T cell polarization to Th1, secreting pro-inflammatory cytokines such as TNF-α, IL-6, IL-1β, IL-12, and type I interferons. Conversely, M2 macrophages are primarily involved in fungal and parasitic infections, tissue remodeling, and repair. The M2a subtype synthesizes cytokines such as IL-4, IL-10, and IL-13, playing a role in allergies, anti-inflammatory activity, and the induction of fibrosis. The M2b subtype produces cytokines such as TNF-α, IL-6, IL-1β, and IL-10, contributing to Th2 polarization and immune regulation. M2c macrophages synthesize IL-10, IL-6, and TNF-α, displaying immunosuppressive, reparative, and remodeling activities. Lastly, the M2d subtype synthesizes IL-10, VEGF (vascular endothelial growth factor), and TGF-β (transforming growth factor beta), playing key roles in angiogenesis and the degradation of cellular debris and apoptotic bodies **[41]**. However, despite MVsplE and MVsplS belonging to the same strain (*L. plantarum*), we observed a pro-inflammatory effect in RAW 264.7 cells stimulated with MVsplS up to hour 5, prior to the challenge, with predominant expression of IL-1β and TNF-α. These differences may be associated with the specific cargo molecules carried during the co-culture of *L. plantarum* with enteropathogenic strains **[14, 16]**. Although further characterization of the cargo molecules in MVsplE compared to MVsplS is required, previous studies have described that the bacterial growth microenvironment influences both the quantity of MVs and the components they carry. In future trials, it would be interesting to evaluate the protein plasticity of MVsplE and MVsplS using 2D electrophoresis, mass spectrometry, and ionic exchange chromatography.

Macrophages stimulated with MVsplS exhibited higher IL-1β expression at hour 3 compared to those stimulated with commercial *E. coli* LPS (Fig 8B). The higher IL-1β expression observed in MVsplS but not in MVsplE could be related to findings by Sandanusova (2024), who demonstrated that the cargo loading process in MVs of *L. rhamnosus* is particular and selective **[14]**. Thus, the co-culture of *L. plantarum* with *S*. Typhimurium could induce the expression and transport of specific antigens in its membrane vesicles (MV), leading to a pro-inflammatory profile in the evaluated cells. In contrast, MVs obtained from the co-culture of *L. plantarum* with *E. coli* induced an anti-inflammatory profile in RAW 264.7 cells.

Additionally, we found that macrophages stimulated with MVsplE or MVsplS triggered a faster and more sustained response to IL-1β and TNF-α than their whole-cell counterparts. Similar results were reported by Briaud (2020) and Toyofuku (2023), who described cargo molecules present in MVs as being more abundant, more concentrated, and mostly biologically active, leading to a more substantial antigenic effect than their whole-cell origins **[13, 18]**.

On the other hand, the cytokine expression kinetics in RAW 264.7 cells stimulated with MVsplE and MVsplS prior to the challenge showed a peak in IL-1β and TNFα expression at hour 3. This contradicts findings by Gutiérrez (2023), who stimulated RAW 264.7 cells with MVs from *L. acidophilus* and observed that the peak expression of IL-1β and TNF-α occurred at hour 1 post-stimulation **[25]**. Furthermore, a similar effect has been observed in OMVs from Gram-negative bacteria. Ávalos (2015) reported that stimulating ovine macrophages with 5 and 10 µg of OMVs from *Mannheimia haemolytica* A2 increased IL-1β and TNFα gene transcription, with the expression peak for both cytokines occurring at hour 1 **[42]**. In both cited studies, a single stimulus was applied to the evaluated cell lines; in contrast, the present study stimulated RAW 264.7 cells at hours 0, 2, and 4. This repeated stimulation could lead to a reinforcement of the response in macrophages, resulting in increased expression of IL-1β and TNF-α upon secondary exposure to the same antigen **[43, 44]**. As the experiment progressed, at hour 5 prior to the challenge, IL-1β and TNFα expression decreased, while IL-10 expression increased, reaching its peak at the end of the challenge (hour 8) with enteropathogenic bacteria.

This phenomenon can be explained by Gharavi (2022), who stated that the inflammatory process begins with pathogen-associated molecular patterns (PAMPs), which activate a signaling cascade that drives macrophages toward an M1 phenotype, characterized by the expression of pro-inflammatory cytokines **[41]**. The next step at the tissue level would involve M1 macrophages clearing cellular debris and initiating repair processes, during which they adopt an anti-inflammatory profile and polarize toward the M2 phenotype. The success of the immune response against various diseases and pathogens depends on the delicate balance between M1/M2 polarization, as both an adequate inflammatory response and subsequent repair are necessary **[45]**. It has been reported that MVs activate macrophages because different PAMPs from the cell of origin are present on their structure and interact with various pattern recognition receptors (PRRs), including TLR2, TLR3, TLR7, TLR9, and NOD2 **[20]**. Once they have been internalized, they exert an effect similar to that of their bacterial origin. Hu et al., in 2020, mention that stimulating RAW 264.7 cells with OMVs of *E. coli* Nissle 1917 improves phagocytosis by increasing the activity of acid phosphatase and iNOS (inducible nitric oxide synthase) **[46]**. In addition, Lee et al. (2021) showed that *L. plantarum* MVs inhibit the growth of *Shewanella putrefaciens* on trypticase soy agar **[28]**. In the other hand the MVs not only improve the phagocytic capacity of macrophages, it has been described that MVs are also capable of inducing their polarization (Kim, et al. 2020) in addition to generating an immunomodulatory response through the expression of IL-10 **[47, 48]**.

Couper (2008) highlighted the crucial role of IL-10 in preventing tissue and/or cellular damage due to a previously exacerbated and uncontrolled immune response. IL-10 can inhibit the expression of pro-inflammatory cytokines, including IL-1β, IL-6, IL-12, IL-18, and TNF-α, in macrophages **[45]**.

Due to this balance between M1/M2 polarization and the action of IL-10, induced by stimulation with MVsplE and MVsplS, we observed a decrease in IL-1β and TNFα expression prior to the challenge with enteropathogenic bacteria. Toward the end of our experiment (hour 8) and during the challenge with enteropathogenic bacteria, IL-10 reached its peak expression in the presence of either *S*. Typhimurium or *E. coli*. In contrast, TNFα expression occurred in parallel with IL-10. This could be related to the balance between the pro-inflammatory and anti-inflammatory responses during the challenge, aiming to prevent an excessive response that could cause irreversible cellular damage **[45]**. Samanta (2021) reported that whole cells of *L. plantarum* isolated from healthy humans can promote anti-inflammatory characteristics during an *S*. Typhimurium infection in Caco-2 cells via IL-10 expression **[49]**. This study showed that the peak IL-10 expression remained unaltered in both *L. plantarum*-treated cells and those co-cultured with *S*. Typhimurium. In the present study, we found that IL-10 expression in macrophages stimulated with MVsplE or MVsplS not only persisted in the presence of enteropathogenic bacteria during the challenge but also increased compared to cells stimulated only with MVs and not exposed to the challenge.

It is essential to note that the genus *Salmonella* can induce macrophage death through IL-1β-mediated pyroptosis **[50, 51]**. However, in our experiment, we observed that IL-1β expression in macrophages remained at basal levels during the challenge with *S.* Typhimurium.

Finally, it is interesting that IL-10 and TNFα expression occurred simultaneously without inhibiting each other. This phenomenon has two possible explanations. The first, as described by Daftarian and Meisel (both in 1996), suggests that following a pro-inflammatory event, TNFα expression induces and enhances IL-10 expression, gradually shifting from a pro-inflammatory to an anti-inflammatory response **[52, 53]**. The second explanation relates to findings by Wang Le (2019), who reported that the M2b macrophage subtype simultaneously produces IL-10 and TNFα, with the latter playing an immunomodulatory role **[54, 55]**. However, further evaluation of markers defining macrophage differentiation from M1 to M2 in culture is needed to fully explain the observations in this experiment.

## Conclusion

This study is the first to report the bactericidal effect of MVs derived from *L. plantarum* isolated from free-living rats against *E. coli* and *S.* Typhimurium, demonstrating stronger antimicrobial activity than C.C. Additionally, MVs stimulated macrophage activation and cytokine expression, promoting an immunomodulatory effect. These findings suggest a promising new approach for preventing and controlling infectious diseases through MVs based on the use of MVs as acellular biologics for alternative treatment.

## Supporting information

**S1 Fig.**
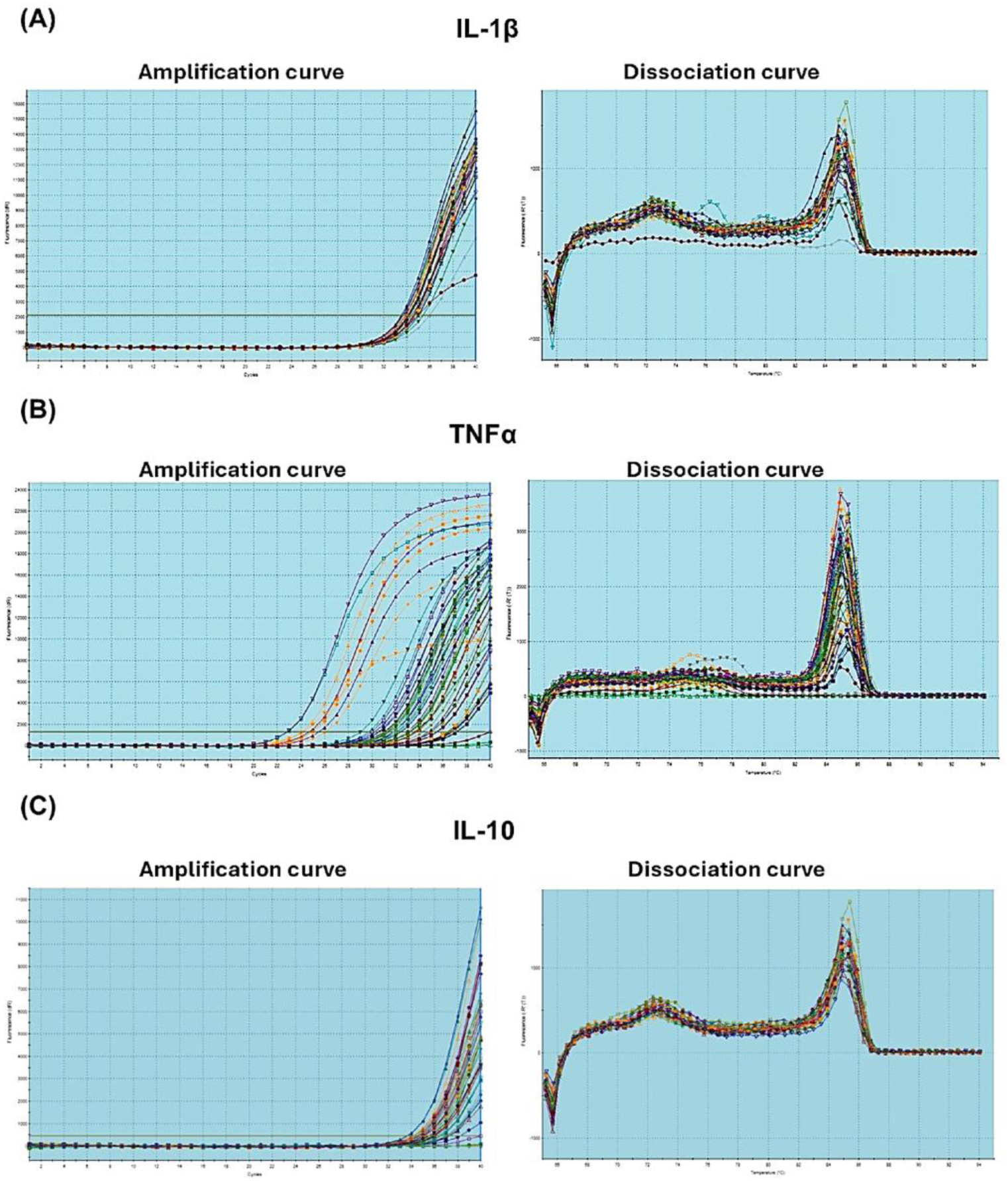
Dissociation and amplification curves for validation and specificity of the primers synthesized for qPCR. (A). IL-1β, (B) TNF α, (C) IL-10.

**S2 Fig.**
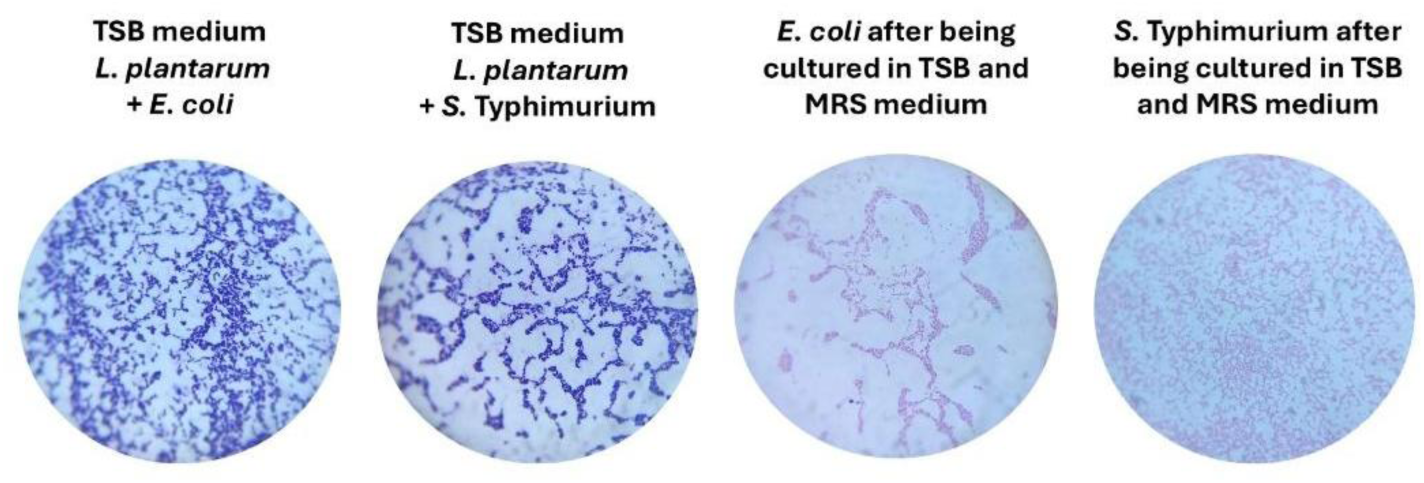
Gram staining of *L. plantarum* co-cultures (1000x). (A) *L. plantarum* strain after being co-cultured with *E. coli* 1×10⁸ in TSB medium and (B) *L. plantarum* strain after being co-cultured with *S*. Typhimurium 1×10⁸ in TSB medium. (C) *E. coli* strain, after being cultured in TSB and MRS, and (D) *S*. Typhimurium strain, after being cultured in TSB and MRS, were used as experimental controls.

**S1 Video.**
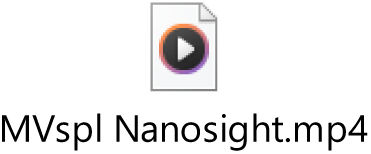
Video from MVspl by NanoSight NS300.

**S2 Video.**
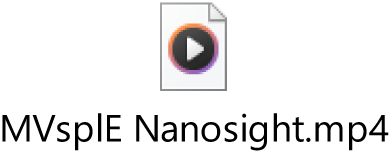
Video from MVsplE by NanoSight NS300.

**S3 Video.**
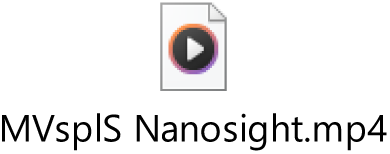
Video from MVsplS by NanoSight NS300.

## Acknowledgments

We thank the biologists. María de Lourdes Rojas Morales from the Advanced Microscopy Laboratory for her support with the transmission electron microscopy micrographs, as well as the Cell Biology Department where the cell cultures were performed, both from to the Centro de Investigación y de Estudios Avanzados del IPN (CINVESTAV).

## Author Contributions

Conceptualization: CGR and CDLS; methodology: HRA, AVR, JLMH, FRGD, MLH; validation: RIHP, HACC and JACO; investigation and resources: CGR; data curation: CDLS.; writing—original draft preparation: GRR, CDLS and CGR; writing—review and editing: GRR, CDLS and CGR; visualization, supervision: GRR, CDLS and CGR, project administration: CGR, and funding acquisition, CGR and JACO. All authors have read and agreed to the published version of the manuscript.” All authors have read and agreed to the published version of the manuscript.”.

## Institutional Review Board Statement

The research project was carried out in accordance with the ethical and humanitarian guidelines that govern experimentation with animals, which are described in NOM-062-ZOO-1999. The animal study protocol was approved by Institutional Subcommittee for the Care and Use of Experimental Animals (SICUAE-FESC) of the Facultad de Estudios Superiores Cuautitlán, UNAM, approval number MC-2022/1-1, dated March 27, 2022.

## Informed Consent Statement

“Not applicable.” for studies not involving humans.

## Data Availability Statement

Data contained within the article.

## Abbreviations

The following abbreviations are used in this manuscript:

BSA: Bovine serum albumin
C.C: Complete cells
C.Cpl: Whole cells of *Lactiplantibacillus plantarum* without co-cultivation
C.CplE: Whole cells of *Lactiplantibacillus plantarum* co-cultured with *E. coli*
C.CplS: Whole cells of *Lactiplantibacillus plantarum* co-cultured with *Salmonella* Typhimurium
CFU: Colony-forming unit
CTAB: Cetyltrimethylammonium bromide
GIT: Gastrointestinal tract
HPRT: Hypoxanthine-guanine phosphoribosyltransferase
iNOS: inducible nitric oxide synthase
IL-1β: Interleukin 1 beta
IL-4: Interleukin 4
IL-6: Interleukin 6
IL-10: Interleukin 10
IL-12: Interleukin 12
IL-13: Interleukin 13
LAB: Lactic Acid Bacteria
LPS: Lipopolysaccharide
MRS: Man–Rogosa–Sharpe
MVs: Membrane vesicles
MVspl: Membrane vesicles of *Lactiplantibacillus plantarum* without co-cultured
MVsplE: Membrane vesicles of *Lactiplantibacillus plantarum* co-cultured with *E. coli*
MVsplS: Membrane vesicles of *Lactiplantibacillus plantarum* co-cultured with *Salmonella* Typhimurium
OMVs: Outer membrane vesicles
PAMPs: Pathogen-associated molecular patterns PRRs Pattern recognition receptor
PBS: Phosphate-buffered saline
SIM: Sulfide, Indole, Motility
TEM: Transmission electron microscopy
TGF-β: Transforming growth factor beta
TLR2: Toll-like Receptor 2
TNF-α: Tumor Necrosis Factor Alpha
TSB: Tryptic Soy Broth
VEGF: vascular endothelial growth factor

